# Stressed alga-free coral cells can initiate coral bleaching

**DOI:** 10.64898/2026.09.30.755578

**Authors:** Jinsong Wei, Yun Bai, Ruijie Wu, Jinru Xie, Jiali Zheng, Miaomiao Luo, Minshuai Zhu, Chenxi Li, Siqi Xie, Kaiyue Han, Kaihui Xu, Minjie Hu

## Abstract

Coral bleaching is driving the rapid decline of coral reefs worldwide, yet a fundamental question remains unresolved: does the bleaching cascade begin in the coral host or in its intracellular algal symbionts, *Symbiodiniaceae*^1^? Resolving where this process starts is critical for understanding how symbiosis collapses and where early intervention might be effective. Here we use transient stress to create a temporal window between the early cellular response and subsequent symbiosis breakdown. Across transient menthol and thermal stress in soft and stony corals, bleaching developed after the stress had ended. In *Xenia* sp.^2^, single-cell transcriptomics revealed a broad host response before detectable symbiont loss, including coordinated redox changes consistent with conditions favouring reactive oxygen species (ROS) accumulation. Cell-resolved imaging showed that alga-free coral cells were the first to become ROS-positive, whereas *Symbiodiniaceae* showed no detectable ROS signal. This early host response was transmissible. Conditioned medium from stressed cultures induced the emergence of ROS-positive host cells and subsequent symbiosis breakdown in unstressed recipients. Purified ROS-positive alga-free coral cells alone were sufficient to trigger symbiosis breakdown in recipient cultures without direct stress exposure. Suppressing the early emergence of ROS-positive coral cells markedly reduced later bleaching, whereas intervention after stress was substantially less effective. These findings identify a host-initiated, intercellularly propagating route to coral bleaching and reveal an early host response that can determine the subsequent fate of the symbiosis.

## Main

The breakdown of the intracellular symbiosis between cnidarian hosts and *Symbiodiniaceae* leads to coral bleaching, a major consequence of marine heat stress and a principal driver of reef decline^3,4^. Without substantial reductions in greenhouse-gas emissions, a majority of coral reefs are projected to experience severe bleaching every year by the mid-2050s^5^, leaving limited time to improve coral resilience. Yet despite decades of research, which partner initiates symbiosis breakdown—the photosynthetic symbiont or the host— remains unresolved^1,6^, making it unclear whether bleaching mitigation should target the host or the symbiont.

A long-standing model places stress-induced dysfunction of *Symbiodiniaceae* at the start of the bleaching cascade. Heat and excess light can impair algal photosynthesis and increase ROS production, leading to the proposal that oxidative stress originating in the symbiont triggers host responses and ultimately destabilizes the symbiosis^7–11^. However, several observations challenge a simple symbiont-initiated model. Bleaching can occur in darkness^12^, and symbiont photodamage or ROS accumulation does not consistently precede symbiont loss^13–16^. On the other hand, Host processes, including programmed cell death, autophagy and vesicular trafficking, have also been implicated in bleaching^17–25^, but it remains unclear whether they help initiate symbiosis breakdown or instead reflect broader responses to stress. Thus, although both partners respond to stress, it remains unclear where the bleaching cascade begins.

Determining who initiates bleaching has been difficult because environmental stress acts simultaneously on the host and symbiont, making their early responses difficult to disentangle, while the continuous progression towards symbiosis breakdown makes initiating events difficult to distinguish from later consequences. Here we use transient stress to create a temporal window between early cellular responses and subsequent symbiosis breakdown, allowing events associated with bleaching initiation to be resolved before detectable symbiont loss. Combining single-cell and functional approaches, we identify a host-initiated route in which an early host response can spread between cells and drive symbiosis breakdown under both menthol and thermal stress.

### Transient menthol exposure elicits a bleaching process

To distinguish early events that initiate bleaching from later changes associated with symbiont loss, we established a transient-stress approach that temporally separates the early cellular response from subsequent symbiont loss. Menthol has been widely used to induce bleaching in corals and sea anemones, typically through repeated or prolonged treatments over days to weeks^26,27^. In *Xenia* sp.^2^, we found that a single 16-h exposure to 0.19 mM menthol was sufficient to initiate the process of bleaching. After 16 h, menthol was removed, and the polyps were washed and maintained in untreated seawater for the remainder of the experiment (Fig. 1a). By day 7, the transiently exposed polyps showed marked paling and tissue contraction, consistent with bleaching (Fig. 1b). Flow-cytometric quantification of alga-hosting cells showed no detectable symbiont loss at the end of the 16-h menthol exposure, but a significant reduction at 88 h, i.e. 72 h after menthol removal (Fig. 1c and Extended Data Fig. 1). Thus, a brief menthol exposure initiated a bleaching process that continued after the stressor had been removed. This temporal separation between stress exposure and detectable symbiont loss provided a defined window in which to resolve events associated with bleaching initiation rather than its later consequences.

**Fig. 1.**
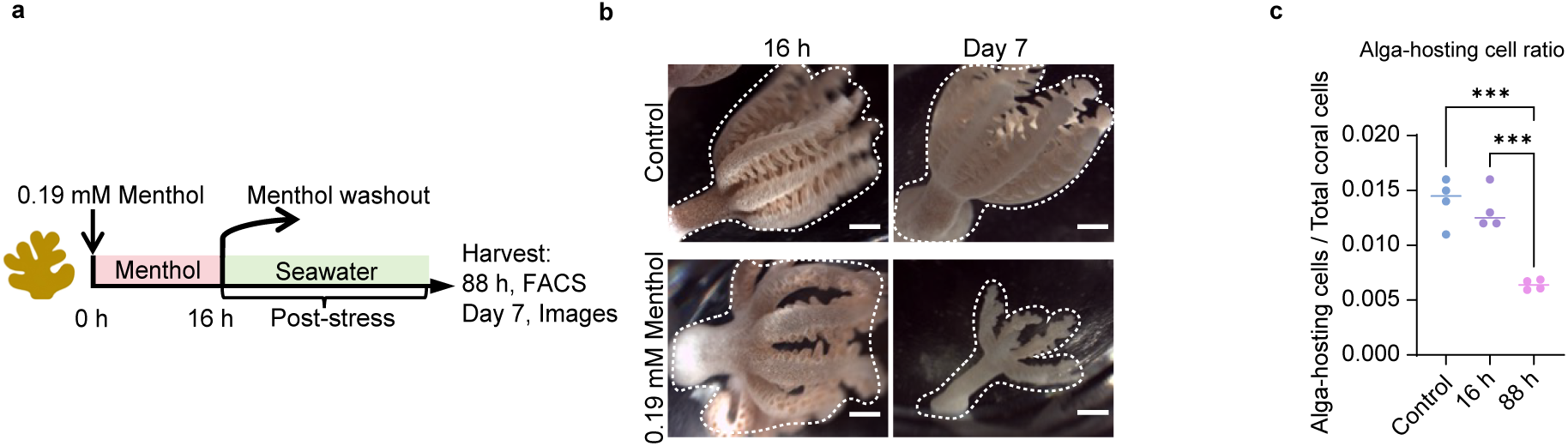
Transient menthol exposure initiates delayed bleaching. **a,** Schematic of the transient menthol-stress experiment. *Xenia* sp. polyps were exposed to 0.19 mM menthol for 16 h, after which menthol was removed and the polyps were maintained in seawater during the post-stress period. **b,** Representative stereomicrographs of control and menthol-treated *Xenia* sp. polyps at 16 h and day 7. Dashed lines outline individual polyps. Scale bars, 0.1 cm. **c,** Proportion of alga-hosting cells among total coral cells in control polyps and in menthol-treated polyps at 16 h and 88 h; Points represent independent polyps and horizontal lines indicate means; *n* = 4. Data are mean ± s.e.m.; one-way ANOVA followed by Dunnett’s test (each group versus 88 h); ns, not significant; ***, *P* < 0.001.

### A broad host transcriptional response precedes symbiont loss

To examine cellular changes before and during symbiont loss, we performed single-cell RNA sequencing of *Xenia* polyps at 16 h, at the end of menthol exposure when alga-hosting cell abundance remained unchanged, and at 88 h, when symbiont loss was evident. We analysed these data together with previously published single-cell profiles from untreated *Xenia* polyps^2^. The combined dataset resolved 37 transcriptional clusters. By SAMap-based^28^ comparison with the *Stylophora pistillata* cell atlas^29^, we identified the major cell types, including endosymbiotic cells, gastrodermal cells, epidermal cells, neurons, and cnidocytes (Fig. 2a and Extended Data Fig. 2a, b).

**Fig. 2.**
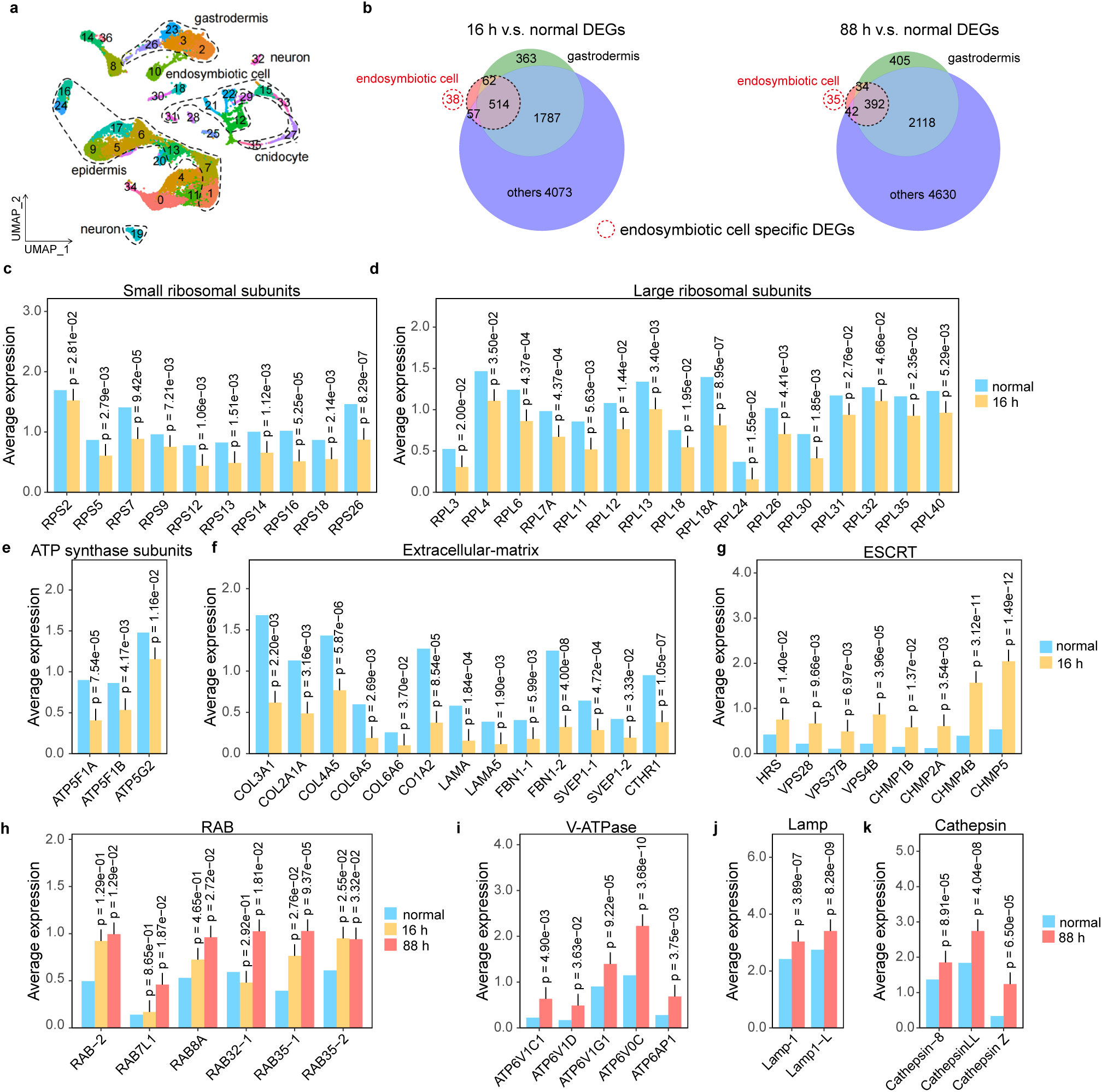
Single-cell transcriptional changes after transient menthol exposure. **a,** UMAP representation of *Xenia* sp. single-cell transcriptomes, resolving 37 transcriptional clusters and the major annotated cell populations. **b,** Overlap of differentially expressed genes (DEGs) in endosymbiotic cells, gastrodermis and other host cell populations at 16 h and 88 h relative to the normal condition. Red dashed circles indicate DEGs specific to endosymbiotic cells. **c–k**, Average expression of Small ribosomal subunits (**c**), Large ribosomal subunits (**d**), Mitochondrial ATP synthase subunits (**e**), Extracellular-matrix genes (**f**), ESCRT components (**g**), RAB-family GTPases (**h**), V-ATPase subunits (**i**), Lamp-family genes (**j**) and Cathepsin-family proteases (**k**) in endosymbiotic cells under the indicated conditions. *P* values are shown above individual genes.

Using memento^30^, we performed differential-expression analysis within each cluster, comparing the 16-h and 88-h conditions separately with the normal condition. Menthol exposure induced broad transcriptional changes across most host cell populations (Extended Data Fig. 2c, d). At 16 h and 88 h, endosymbiotic cells showed a total of 671 and 503 differentially expressed genes, respectively, of which only 38 and 35 were unique to endosymbiotic cells. Most transcriptional changes were shared with other host populations, including both gastrodermal and non-gastrodermal cells. (Fig. 2b and Supplementary Table 1, 2). The few endosymbiotic-cell-specific genes were dominated by established markers of these cells and did not reveal an obvious functional programme associated with known bleaching processes (Supplementary Table 3, 4). If the early bleaching response were driven primarily by symbiont dysfunction, endosymbiotic cells, which are in direct contact with *Symbiodiniaceae*, would be expected to show a more distinct transcriptional response than other host cell populations. Instead, the limited number of endosymbiotic-cell-specific changes, together with the broad response across other host populations, indicates that the early bleaching response is unlikely to be initiated primarily by intracellular *Symbiodiniaceae*.

We next examined transcriptional changes in endosymbiotic cells at 16 h. Genes encoding both small and large ribosomal subunits were broadly downregulated (Fig. 2c, d and Extended Data Fig. 3a,b), consistent with a stress-associated reduction in protein synthesis^31^. Transcripts encoding mitochondrial ATP synthase subunits, including ATP5F1A, ATP5F1B and ATP5G2, were also reduced (Fig. 2e and Extended Data Fig. 3c), suggesting reduced mitochondrial ATP-generating capacity^32^. Several extracellular-matrix genes declined in parallel, including multiple collagen genes, LAMA5, FBN1 and SVEP1 paralogues, and CTHR1 (Fig. 2f and Extended Data Fig. 3d). The reduction in extracellular-matrix transcripts is consistent with previous observations of host-cell detachment during cnidarian bleaching^25,33^. Many of these early transcriptional changes were transient, with differential expression either weakening or shifting in the opposite direction by 88 h (Extended Data Fig. 3 and Supplementary Table 5).

Endomembrane trafficking was also progressively remodelled at the transcriptional level. At 16 h, multiple components of the ESCRT machinery were upregulated, including HRS, VPS28, VPS37B, VPS4B and several CHMP proteins(Fig. 2g), spanning cargo recognition, endosomal sorting, membrane scission and ESCRT recycling^34^. RAB35 paralogues, which are involved in endocytic recycling and plasma-membrane trafficking^35^, were also increased at this stage (Fig. 2h). Together, these changes indicate coordinated remodelling of endosomal sorting and membrane-recycling pathways before detectable symbiont loss. By 88 h, the response had expanded to additional RAB-family GTPases, including RAB2, RAB7L1, RAB8A and RAB32 (Fig. 2h and Extended Data Fig. 3e), indicating broader remodelling of intracellular membrane-trafficking pathways^36^. In parallel, V-ATPase components, LAMP1-family proteins and cathepsin-family proteases were upregulated (Fig. 2i–k and Extended Data Fig. 3f–h). These genes encode machinery associated with endolysosomal acidification, lysosomal membrane function and proteolysis^37,38^, respectively, indicating increasing transcriptional engagement of lysosome-associated processes by 88 h. Together, these expression changes indicate a temporal progression from early remodelling of endosomal sorting and membrane-recycling machinery at 16 h to broader engagement of endolysosomal and lysosome-associated programmes by 88 h. Such changes are compatible with cellular routes of symbiont loss described during cnidarian bleaching: altered membrane-trafficking pathways could facilitate symbiont expulsion, whereas increased expression of endolysosomal and lysosomal machinery is consistent with intracellular degradation pathways such as symbiophagy^24–26^.

Many of these transcriptional changes were also detected in other host cell populations, with their magnitude and distribution varying among cell types (Extended Data Fig. 3).

Together, these data place widespread host transcriptional changes before detectable symbiont loss at 16 h and reveal progressive endomembrane and lysosome-associated changes as bleaching develops.

### ROS first appear in alga-free coral cells

Because ROS have long been implicated in coral bleaching^7–11,39^, we examined redox-associated transcriptional changes. At 16 h, in endosymbiotic cells, G6PD, which supplies NADPH to glutathione- and thioredoxin-dependent antioxidant systems^40^, was reduced.

SOD1, which converts superoxide to hydrogen peroxide and oxygen^41^, was also downregulated, together with PRDX5 and PRDX6, peroxiredoxins with peroxide-reducing activity^42^ (Fig. 3a). By contrast, NOX5, a calcium-regulated NADPH oxidase that generates superoxide^43^, and CYP1B1, whose cytochrome P450 activity can generate ROS under uncoupled conditions^44^, were upregulated (Fig. 3b). At the same time, several genes that could counteract oxidative and proteotoxic damage were upregulated. GLRX and TXNRD3 support thiol redox homeostasis by restoring oxidized protein thiols and maintaining thioredoxin-dependent reducing capacity^45,46^, respectively. Ferritin-like genes can sequester labile iron, thereby limiting Fenton chemistry and the formation of highly reactive oxygen species^47^. Several heat-shock proteins were also induced (Fig. 3c). Although they do not directly remove ROS, these molecular chaperones can buffer the downstream consequences of oxidative stress by stabilizing and refolding damaged proteins^48^. Many of these redox-associated transcriptional changes were also observed across other host cell populations (Extended Data Fig. 4 and Supplementary Table 6). Together, these transcriptional changes indicate coordinated remodelling of host redox regulation, with changes that could favour ROS accumulation occurring alongside increased expression of genes that can limit oxidative and cellular damage at 16 h.

**Fig. 3.**
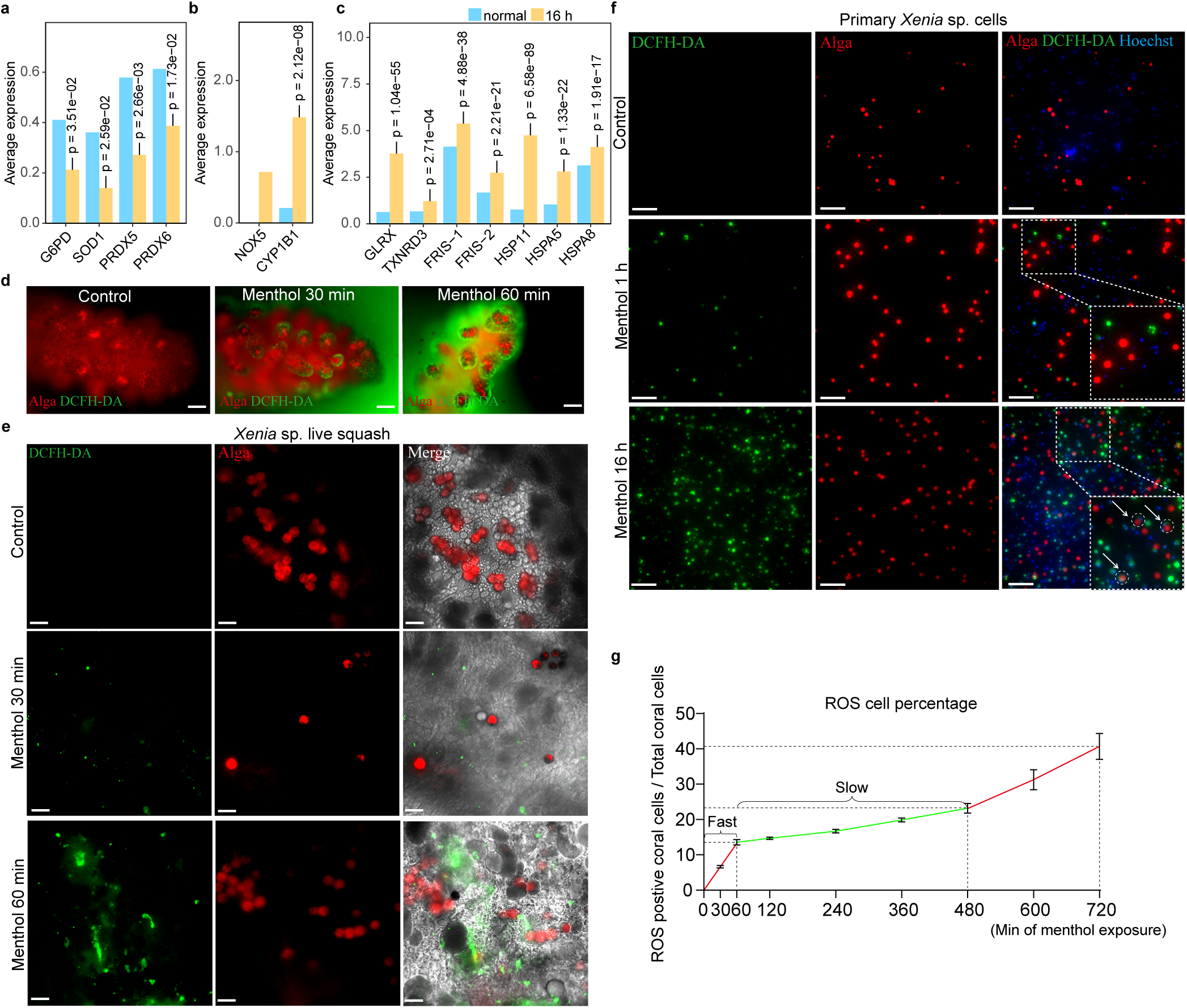
Host-cell ROS emerge before symbiont loss. **a–c,** Average expression of G6PD, SOD1, PRDX5 and PRDX6 (**a**), CYP1B1 and NOX5 (**b**), and GLRX, TXNRD3, ferritin-like genes and heat-shock proteins (**c**) in endosymbiotic cells under the indicated conditions. *P* values are shown above individual genes and were obtained from differential-expression analysis using Memento. No *P* value is shown for the gene with no detectable expression in one of the compared groups. **d,** Representative images of DCFH-DA fluorescence in intact live *Xenia* sp. polyps before menthol exposure and after 30 and 60 min of exposure. Algae autofluorescence is shown in red and DCFH-DA fluorescence in green. Scale bars, 100 µm. **e,** Representative images of live squash preparations of *Xenia* sp. before menthol exposure and after 30 and 60 min. Scale bars, 50 µm. **f,** Representative images of *Xenia* sp. primary-cell cultures before menthol exposure and after 1 h and 16 h of menthol exposure. DCFH-DA fluorescence is shown in green, algal autofluorescence in red and Hoechst staining in blue. Insets show enlarged regions of the merged images. Arrows indicate alga-hosting cells. Scale bars, 100 µm. **g,** Time course of the proportion of ROS-positive host cells during 720 min of menthol exposure. ROS-positive cells increased rapidly during the first 60 min, accumulated more slowly thereafter and increased more rapidly again at later time points. Data are mean ± s.e.m.; *n* = 3 independent cultures.

Prompted by this transcriptional pattern, we used the ROS-sensitive fluorescent probe DCFH-DA to determine where ROS signals first appeared during menthol exposure. Within 30–60 min, DCFH-DA-positive cells were first detected near the tentacle tips, after which the signal spread more broadly across the tissue and extended into the gastrodermis by 16 h (Fig. 3d and Extended Data Fig. 5). At the early time points, the DCFH-DA signal was detected predominantly in alga-free host cells (Fig. 3e). The *Symbiodiniaceae* showed no detectable signal throughout the menthol exposure (Fig. 3d, e and Extended Data Fig. 5). Because alga-hosting cells and *Symbiodiniaceae* lie within the gastrodermis, their later ROS signal could reflect reduced or delayed exposure to menthol rather than an intrinsic difference in stress responses between cell populations.

To address this possibility, we established primary-cell cultures of *Xenia* (Extended Data Fig. 6a, b). This allowed different host cell populations to be exposed directly to the same extracellular menthol concentration. We treated the primary-cell cultures with a similar menthol stress (10 µM) (Fig. 3f), ROS-positive cells again first appeared among alga-free host cells, whereas alga-hosting cells became ROS-positive only after longer exposure (Fig. 3f). Both intracellular and free-living *Symbiodiniaceae* showed no detectable ROS signal throughout the assay. ROS-positive host cells accumulated rapidly during the first 60 min, increased more slowly from 60 to 480 min, and then rose slightly sharper again between 480 and 720 min (Fig. 3g and Extended Data Fig. 6c, d).

### ROS-positive host cells drive symbiosis breakdown

The progressive spread of ROS-positive cells across the tissue and primary-cell cultures raised the possibility that the early host response could propagate from initially responding cells to other cells. We therefore tested whether an extracellular activity released from stressed cells could transmit this response to unstressed cells. Donor *Xenia* primary cultures were exposed to menthol for 2 h, washed, and then incubated in fresh medium for a further 4 h. Conditioned medium was collected only during this post-treatment period and was transferred to untreated recipient cultures that had never been exposed to menthol. A cell-free control processed in parallel through the same menthol exposure and medium-replacement steps controlled for residual menthol carryover, whereas conditioned medium from untreated donor cultures controlled for factors released under normal culture conditions (Fig. 4a). ROS-positive cells were detected in recipient cultures within 4 h of conditioned-medium transfer and became more abundant by 16 h (Fig. 4b). Flow-cytometric quantification showed that the proportion of ROS-positive host cells increased between 4 and 16 h, reaching approximately 40% at 16 h, whereas both control groups remained low (Fig. 4c and Extended Data Fig. 7). This response was followed by a significant reduction in the proportion of alga-hosting cells at 48 h relative to both controls (Fig. 4d). Thus, transiently stressed donor cultures released a transferable extracellular activity that induced an oxidative response in previously unstressed cells and subsequently drove symbiosis breakdown.

**Fig. 4.**
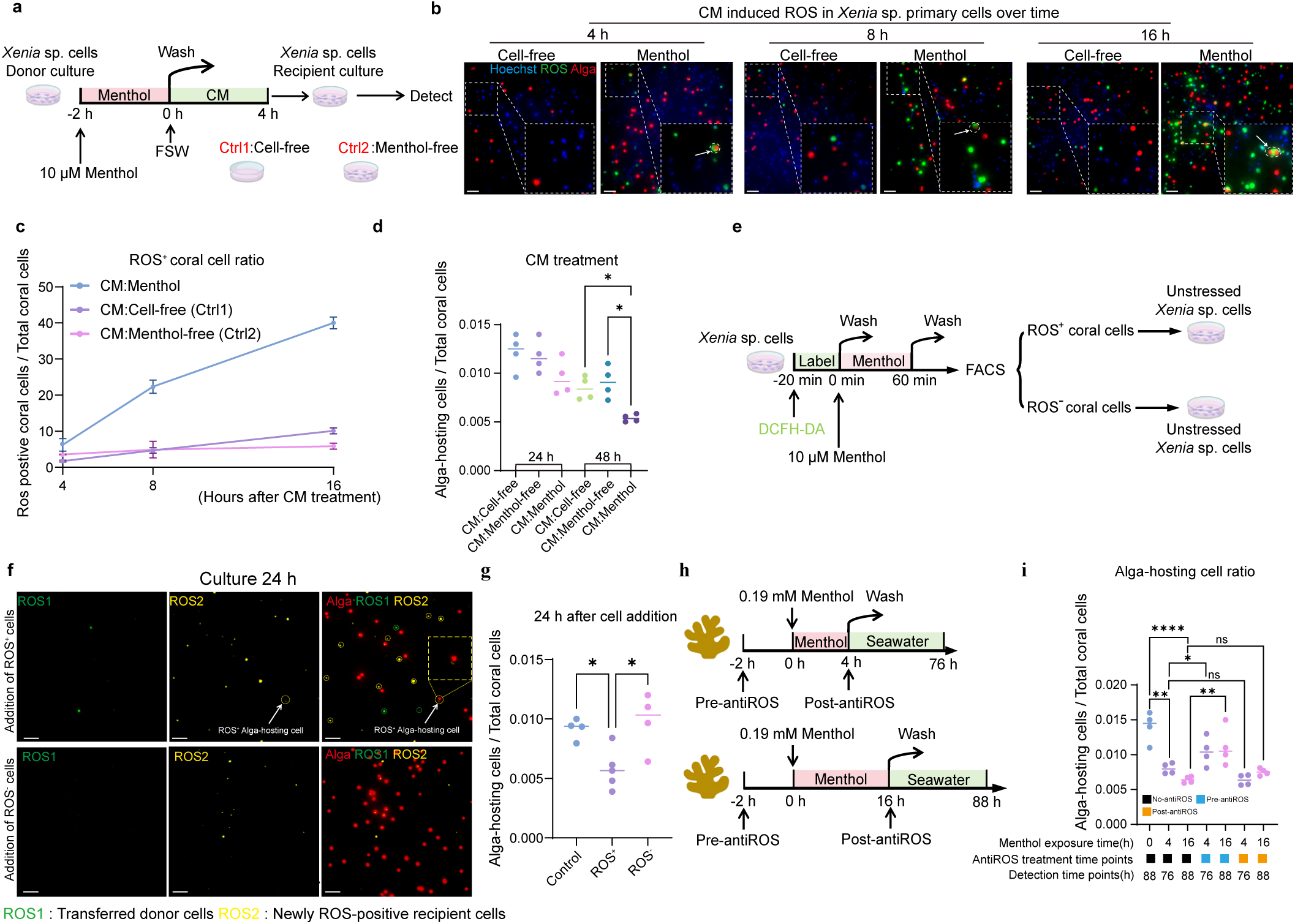
ROS-positive host cells propagate the response and drive symbiosis breakdown. **a,** Schematic of the conditioned-medium (CM) transfer experiment. *Xenia* sp. donor primary cultures were exposed to 10 µM menthol for 2 h, washed, and incubated in fresh medium for a further 4 h to generate conditioned medium, which was transferred to recipient cultures that had not been exposed to menthol. A cell-free control (Ctrl1) subjected to the same menthol-exposure and washout procedure controlled for menthol carryover, and conditioned medium from untreated donor cultures served as a control (Ctr2) for constitutive factors released by cultured cells. FSW, filtered seawater. **b,** Representative recipient cultures at 4, 8 and 16 h after CM transfer, showing the progressive emergence of DCFH-DA-positive host cells. **c,** Proportion of ROS-positive host cells at 4, 8 and 16 h after transfer of Menthol-stressed CM, Cell-free control medium or Menthol-free control CM. **d,** Proportion of alga-hosting cells in recipient cultures at 24 and 48 h after CM transfer. **e,** Schematic of the ROS-positive host-cell transfer experiment. *Xenia* sp. primary cells were labelled with DCFH-DA and exposed to 10 µM menthol for 60 min. ROS-positive and ROS-negative symbiont-free host cells were isolated by fluorescence-activated cell sorting and transferred to recipient cultures that had not been directly exposed to menthol. **f,** Representative images of recipient cultures 24 h after addition of ROS-positive or ROS-negative donor cells. ROS1 marks transferred donor cells, whereas ROS2 marks newly arising ROS-sensitive fluorescence after transfer; algal autofluorescence is shown in red. **g,** Proportion of alga-hosting cells 24 h after addition of ROS-positive or ROS-negative donor cells. **h,** Schematics of the antioxidant-intervention experiments using 16-h or 4-h menthol exposure. Vitamin C treatment was initiated either 2 h before menthol exposure and maintained during stress (pre-antiROS) or after menthol removal (post-antiROS). **i,** Proportion of alga-hosting cells following 4-h or 16-h menthol exposure with no antioxidant, pre-antiROS or post-antiROS treatment. Samples were analysed at 76 h after initiation of the 4-h exposure and at 88 h after initiation of the 16-h exposure. Points represent independent cultures and horizontal lines indicate means. Scale bars, 100 µm. Data are mean ± s.e.m.; n = 4 independent cultures; one-way ANOVA followed by Dunnett’s multiple-comparison test; ns, not significant; *, P < 0.05, **,P < 0.01, ***, P < 0.001, ****, P < 0.0001.

We next tested whether early ROS-positive alga-free host cells that emerged during menthol exposure were sufficient to drive symbiosis breakdown. Primary *Xenia* cultures were stained with DCFH-DA and exposed to menthol for 60 min. ROS-positive and ROS-negative alga-free host cells were then isolated by fluorescence-activated cell sorting and added in equal numbers to untreated recipient cultures that had never been exposed to menthol (Fig. 4e). Transferred donor cells retained the original DCFH-DA-derived signal (ROS-1), whereas newly arising oxidative signals after transfer were detected with a second ROS probe, Dihydroethidium (DHE), in an independent fluorescence channel (ROS-2). Cells positive for ROS-2 but lacking ROS-1 could therefore be distinguished from the transferred donor population. Following transfer of ROS-positive donor cells, ROS-2-positive/ROS-1-negative cells appeared in the recipient cultures (Fig. 4f), indicating that the oxidative response had extended beyond the transferred donor cells. At 24 h after cell addition, cultures receiving ROS-positive donor cells showed a significant reduction in the proportion of alga-hosting cells compared with both untreated controls and cultures receiving matched ROS-negative host cells (Fig. 4g). Thus, transferred ROS-positive host cells were sufficient to propagate the host response and drive symbiosis breakdown in recipient cultures, despite the absence of direct stress exposure.

We therefore tested whether suppressing the emergence of ROS-positive host cells during the early stress phase could reduce subsequent bleaching. We treated polyps with antioxidant, Vitamin C, beginning 2 h before menthol exposure and maintained the treatment throughout the stress period, using both 4-h and 16-h menthol exposures (Fig. 4h). Under both conditions, early Vitamin C treatment markedly preserved the proportion of alga-hosting cells, whereas treatment initiated after menthol removal didn’t show a significant effect (Fig. 4i). Together, these results show that limiting the early emergence of ROS-positive host cells can reduce subsequent bleaching under menthol stress.

### Host-initiated bleaching extends across corals and thermal stress

Having established that an early host response can spread between cells and drive symbiosis breakdown in *Xenia*, we tested whether the same pattern occurred in other corals.

We examined the stony coral *Pocillopora damicornis* and a second *Xenia* species (*Xenia* sp.2) (Extended Data Fig. 8a) with the same 16-h menthol exposure. ROS again first appeared in coral cells near the tentacle tips, whereas most *Symbiodiniaceae* showed no detectable ROS signal during the early response (Fig. 5a, b). In both species, the proportion of alga-hosting cells remained similar to controls at the end of the 16-h exposure but was significantly reduced by 88 h (Fig. 5c, d). Vitamin C treatment initiated 2 h before stress and maintained throughout the exposure markedly reduced subsequent bleaching, whereas treatment initiated after the stress didn’t show a significant effect (Fig. 5c, d). Thus, the early host ROS response and the preferential benefit of intervention during this early phase were observed in both soft and stony corals.

**Fig. 5.**
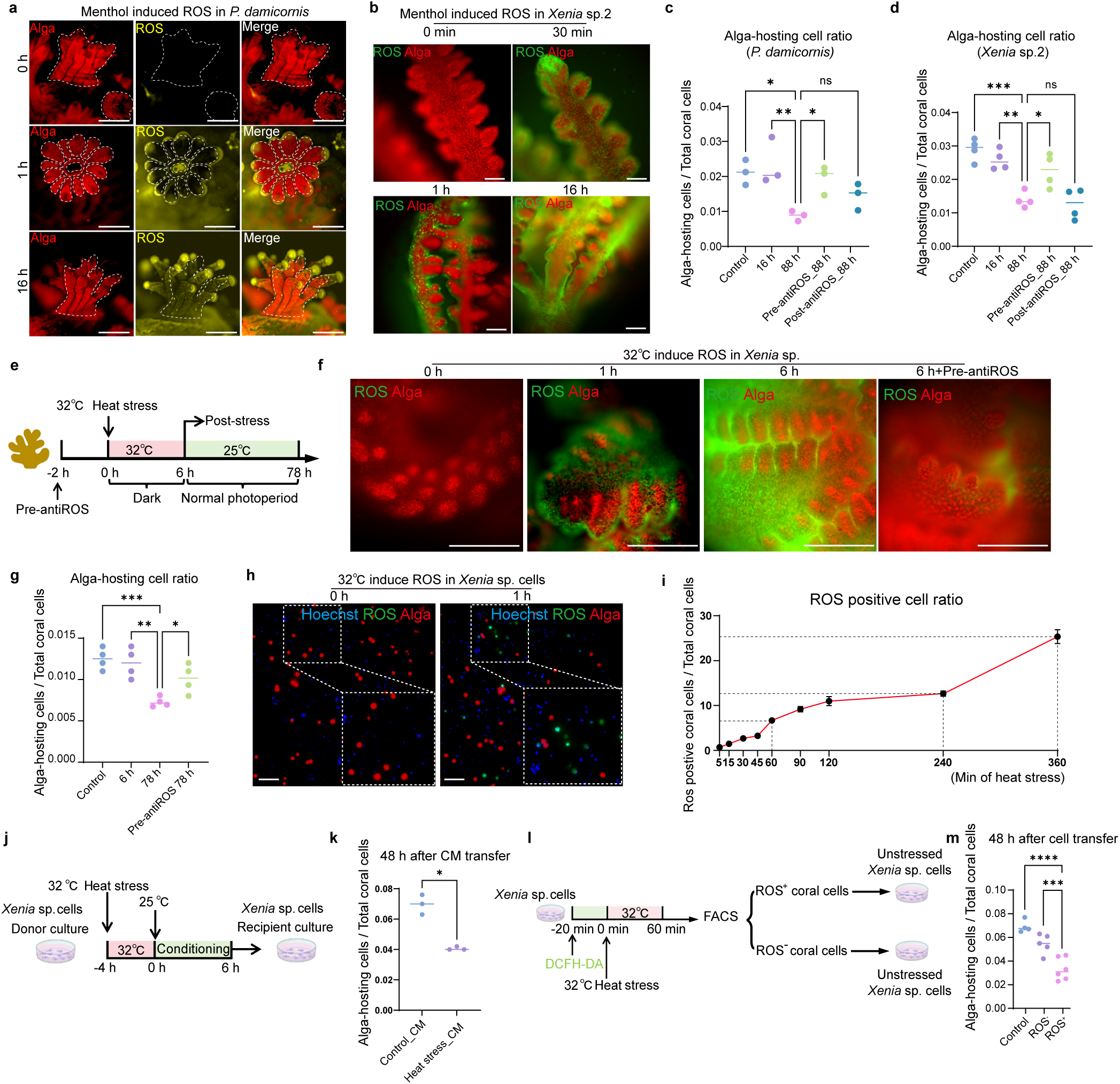
Host-initiated bleaching extends across corals and thermal stress. **a,** Representative images of menthol-induced DHE fluorescence in *Pocillopora damicornis* before treatment and after 1 h and 16 h of menthol exposure. Algae autofluorescence is shown in red and DHE fluorescence in yellow. Scale bars, 0.2 cm. **b,** Representative images of menthol-induced DCFH-DA fluorescence in a second *Xenia* species (*Xenia* sp.2) before treatment and at 30 min, 1 h and 16 h of menthol exposure. Scale bars, 0.1 cm. **c, d,** Proportion of alga-hosting cells in *P. damicornis* (**c**) and *Xenia* sp.2 (**d**) in controls, at 16 h and 88 h after the start of menthol exposure, and at 88 h following pre-stress (pre-antiROS) or post-stress (post-antiROS) vitamin C treatment. **e,** Schematic of the transient heat-stress. *Xenia* polyps were exposed to 32 °C for 6 h in darkness and subsequently maintained at 25 °C under the normal photoperiod until 78 h. In the pre-stress antioxidant group (pre-antiROS), vitamin C treatment began 2 h before heat exposure and was maintained during the stress period. **f,** Representative DCFH-DA fluorescence in *Xenia* sp. polyps before heat stress, after 1 h and 6 h at 32 °C, and after 6 h of heat stress with pre-stress vitamin C treatment. Algal autofluorescence is shown in red and DCFH-DA fluorescence in green. Scale bars, 0.1 cm. **g,** Proportion of alga-hosting cells in control *Xenia* sp. polyps, after 6 h of heat stress, at 78 h after stress initiation, and at 78 h following pre-stress vitamin C treatment. **h,** Representative *Xenia* sp. primary-cell cultures before and after 1 h of exposure to 32 °C, showing DCFH-DA fluorescence in green, algal autofluorescence in red and Hoechst staining in blue. Scale bars, 100 µm. **i,** Time course of the proportion of ROS-positive *Xenia* sp. host cells during 360 min of heat stress at 32 °C. Data are mean ± s.e.m. **j,** Schematic of the heat-stress-conditioned-medium experiment. Donor primary cultures were exposed to 32 °C for 4 h, returned to 25 °C and incubated for a further 6 h to generate conditioned medium, which was transferred to untreated recipient cultures. **k,** Proportion of alga-hosting cells 48 h after transfer of conditioned medium from control or heat-stressed donor cultures. **l,** Schematic of the heat-stressed ROS-positive host-cell transfer experiment. DCFH-DA-labelled *Xenia* sp. primary cells were exposed to 32 °C for 60 min, after which ROS-positive and ROS-negative cells were isolated by fluorescence-activated cell sorting and transferred to recipient cultures that had not been directly exposed to heat stress. **m,** Proportion of alga-hosting cells 48 h after addition of control, ROS-negative or ROS-positive donor cells. Points represent independent polyps (**c, d, g**) or independent primary cultures (**k, m**), and horizontal lines indicate means. Data are mean ± s.e.m.; *n* = 4 independent cultures; one-way ANOVA followed by Dunnett’s multiple-comparison test; ns, not significant; *, *P* < 0.05, **, *P* < 0.01, ***,*P* < 0.001, \*\*\*\**P* < 0.0001.

We then tested whether a similar host-driven bleaching occurred under thermal stress. *Xenia* polyps were exposed to 32 °C for 6 h, mimicking the short-term high-temperature excursions that occur naturally in shallow reef habitats^49,50^, and were kept in darkness to minimize potential concurrent light-dependent photochemical stress (Fig. 5e). During this heat exposure, ROS again first appeared in a subset of alga-free host cells, whereas most *Symbiodiniaceae* showed no detectable ROS signal. The ROS signal subsequently spread to broader regions of the host tissue (Fig. 5f). At the end of the 6-h heat exposure, the proportion of alga-hosting cells remained similar to controls, but declined during the subsequent post-stress period after the polyps had returned to the normal temperature (Fig. 5g). Thus, as under menthol stress, the early oxidative response in host preceded detectable symbiosis breakdown. Vitamin C treatment initiated before and maintained throughout the heat exposure again markedly reduced emergence of ROS-positive host cells and subsequent bleaching (Fig. 5f, g). The same early host response and antioxidant protection were observed in *P. damicornis* and the *Xenia* sp.2 under transient thermal stress (Extended Data Fig. 8b, c).

To test whether the propagating host response observed under menthol stress also operated under thermal stress, we turned to *Xenia* primary-cell cultures, where host cells and *Symbiodiniaceae* experienced the same heat exposure. ROS accumulated in host cells, whereas no detectable ROS signal was observed in *Symbiodiniaceae* within 4 h of 32 °C stress (Fig. 5h, i and Extended Data Fig. 9a, b). Conditioned medium collected after transient heat stress induced an oxidative response in unstressed recipient cultures (Fig. 5j and Extended Data Fig 9c). By 48 h, the proportion of alga-hosting cells was significantly reduced compared with recipients of control conditioned medium (Fig. 5k). Thus, heat-stressed host cells released extracellular activity that could promote symbiosis breakdown without direct heat exposure. We then isolated ROS-positive and ROS-negative host cells after transient heat exposure and transferred equal numbers of each population into unstressed recipient cultures (Fig. 5l. The ROS appeared in the recipient cultures when adding the ROS-positive cell (Extended Data Fig. 9d). By 48-h, cultures receiving ROS-positive host cells again showed a marked reduction in alga-hosting cells compared with control cultures and those receiving matched ROS-negative cells (Fig. 5m). Thus, as under menthol stress, thermal stress generated a host-cell state that could propagate to unstressed cells and promote subsequent symbiosis breakdown.

Together, these experiments establish a common host-initiated sequence across stressors and coral species, in which ROS-positive alga-free host cells emerge before detectable symbiont loss and the host response can propagate to unstressed cells, ultimately leading to symbiosis breakdown. Antioxidant intervention during the early stress phase was able to markedly reduce subsequent bleaching.

## Discussion

A long-standing question in coral bleaching research is whether bleaching is initiated by the host or by its algal symbionts^1^. Our transient-stress experiments resulted in delayed bleaching after the stress had ended, separating the early response to stress from subsequent symbiosis breakdown. Conditioned medium and ROS-positive host cells from stressed cultures both induced ROS responses and drove symbiosis breakdown in unstressed recipient cultures, showing that the response can propagate beyond the initially stressed cells. Together, these results show that the host can drive bleaching under the stresses tested here. Stressors acting primarily on *Symbiodiniaceae* may trigger bleaching through different mechanisms.

The mechanisms linking the early host response to symbiont loss remain unclear. Key questions include which host cells respond first, what extracellular signals spread the response, and how alga-hosting cells receive these signals and trigger symbiosis breakdown. The *Xenia* primary-cell culture system established here provides a tractable platform for resolving these steps and identifying the underlying pathways.

The ability of a transient stress to initiate bleaching may be particularly relevant to bleaching during natural episodes of elevated seawater temperature. Cumulative thermal exposure remains a strong predictor of bleaching risk across reefs^51,52^. Our results, however, show that a brief period of heat stress can be sufficient to initiate bleaching. Such short periods of high temperature also occur naturally on coral reefs^49,50^. Previous experiments have further shown that acute thermal exposure can rapidly alter coral physiology and gene expression^53,54^. Together, these observations raise the possibility that brief periods of high temperature above local tolerance thresholds can act as initiating events for bleaching, with symbiosis breakdown continuing after the acute heat stress has passed. Thus, in addition to cumulative thermal exposure, acute heat events may also need to be considered when predicting bleaching risk.

The delay between heat exposure and detectable bleaching also points to a potential window for bleaching mitigation. Bleaching-level heat stress can now be forecast weeks to months in advance^55^, providing an opportunity to act before bleaching occurs. In our experiments, antioxidant treatment beginning before stress and continuing throughout heat exposure strongly reduced subsequent bleaching. These findings suggest that the early host response could be targeted during this pre-bleaching window, complementing current efforts to improve coral resilience through thermally tolerant corals and *Symbiodiniaceae*^56–58^.

## Acknowledgements

We thank Jinrong Peng, Jun Chen and Mehdi Bouhaddou for discussion and comments on the manuscript, and Shuang Chen for coral husbandry. This work was supported by Zhejiang Provincial Natural Science Foundation of China under Grant No. LR25D060002 (M.H.), National Science Foundation of China under Grant No. 32400083 (M.H.), Fundamental Research Funds for the Central Universities (M.H.), and start-up funds from Zhejiang University (M.H.).

## Author contribution

Y.B., and M.H. conceived and supervised the project. J.W., Y.B., and M.H. designed experiments. J.W., Y.B., J.X., J.Z., M.L., C.L., M.Z., S.X., K.H. and K.X. performed the experiments. R.W. C.L. analysed the data., J.W., Y.B., and M.H. interpreted the data and wrote the manuscript.

